# Ectopic *hAMH*-driven SOX17 expression induces hyperplastic Sertoli valve formation in mouse testes

**DOI:** 10.64898/2026.05.08.723552

**Authors:** Xiao Han, Aya Uchida, Seohyeon Lee, Kosuke Nakamura, Katsuki Takahashi, Tsutomu Endo, Ayaka Yanagida, Ryuji Hiramatsu, Akihiko Kudo, Masami Kanai-Azuma, Yoshiakira Kanai

## Abstract

In the terminal segment of the seminiferous tubules, SOX17 expression in the rete testis (RT) epithelium plays a crucial role in the formation of the Sertoli valve (SV), as revealed by phenotypic analyses of RT-specific *Sox17* conditional knockout (cKO) mouse testes. In these RT-specific *Sox17* cKO testes, SV disruption leads to the backflow of RT fluid into the seminiferous tubules, resulting in defective spermiogenesis and male infertility. Although valve deformation in the *Sox17* cKO testes is likely caused indirectly by impaired downstream actions of *Sox17* in the RT, the mechanisms by which SOX17 in RT influences SV formation in the seminiferous tubules remain unclear. To address this, we generated a novel *AMH-Sox17* transgenic (Tg) mouse line carrying a human *AMH* promoter-driven *Sox17* cDNA cassette. We analyzed the phenotypes of the Sertoli valve and spermatogenesis in *AMH-Sox17* Tg mice, as well as in RT-specific *Sox17* cKO; *AMH-Sox17* Tg double mutant mice. Ectopic SOX17 (SOX17^+^) expression in Sertoli cells resulted in excessive Sertoli valve structures with acetylated tubulin bundles in the terminal segment of the *AMH-Sox17* Tg testes, along with enhanced WNT4/RSPO1 signaling, suggesting the enhanced valve formation of ectopic SOX17^+^ Sertoli cells by themselves. Moreover, the *AMH-Sox17* Tg could partially rescue the SV deformation and infertility in RT-specific *Sox17* cKO mice, leading to proper SV formation, normal spermiogenesis and a partial recovery of male fertility in *AMH-Sox17* Tg; RT-specific *Sox17* cKO double mutant mice. These findings genetically demonstrate that ectopic SOX17^+^ Sertoli cells can compensate for SOX17 paracrine signaling in the RT, underscoring a key shared downstream pathway between RT and SV.

**Summary statement:** The paracrine actions downstream of ectopic SOX17 expression in the Sertoli cells not only promote the valve formation, but also partially rescue the defective spermiogenesis of the rete testis-specific *Sox17*-null mice.

## Introduction

Valves are critical anatomical structures that ensure unidirectional fluid flow in the circulatory and lymphatic systems. They function passively to prevent backflow, maintaining efficient fluid dynamics within vessels (Moore and Bertram, 2018; O’Donnell and Yutzey, 2020). When valve integrity is compromised, it distorts the luminal architecture, reduces intravascular pressure, and may result in regurgitation.

In the mammalian testis — an organ characterized by active, oscillatory fluid flow (Fleck et al., 2021)— a valve-like structure called the Sertoli valve (SV) emerges at the termini of seminiferous tubules (STs) during puberty (Aiyama et al., 2015; Nagasawa et al., 2018; Imura-Kishi et al., 2021). The SV comprises a specialized subset of Sertoli cells that exhibit elevated levels of phosphorylated AKT and STAT3. These cells extend cytoplasmic projections enriched with acetylated tubulin-positive (ace-TUB⁺) microtubules into the rete testis (RT) lumen, forming a dynamic and structurally distinct valve-like barrier (Russell, 1993). Recently, we identified SOX17, a transcription factor expressed in RT epithelial cells, as essential for SV formation and for spermiogenesis in the ST. In mice with RT-specific conditional knockout of *Sox17* (*Sox17* cKO), the SV fails to form properly before puberty, allowing retrograde fluid flow from the RT into the convoluted ST (Uchida et al., 2022). Dysfunction of the RT–SV valve disrupts spermatogenesis in the upstream seminiferous tubules, leading initially to spermatid sloughing with preservation of the Sertoli cell architecture, followed by the progressive involvement of individual seminiferous tubules with age (Uchida et al., 2022; see Fig. S1). These findings imply the critical role of the RT-SV structure in maintaining proper fluid dynamics essential for spermatogenesis.

Interestingly, the SV epithelium provides a specialized niche for undifferentiated spermatogonia (A_undiff_) and immature/proliferative Sertoli cells (Aiyama et al., 2015; see reviews by Figueiredo et al., 2021; Seco-Rovira et al., 2025). The basal compartment of the SV epithelium supports self-renewal of A_undiff_ spermatogonia by maintaining high expression of growth factors such as FGFs (Takashima et al., 2015; Kitadate et al., 2019; Yang et al., 2021; also see the review by Hofmann and McBeath, 2022) and GDNF (Meng et al., 2000; Uchida et al., 2016), together with by locally suppressing retinoic acid (RA)-induced differentiation (Imura-Kishi et al., 2021). Furthermore, SV Sertoli cells display Cyclin D1-positive states in mice and hamsters, and in rats and hamsters, some of these cells proliferate and contribute to the convoluted ST even at adult stage (Aiyama et al., 2015; Figueiredo et al., 2019), suggesting that the SV acts as a niche for proliferative Sertoli cells in hamsters and rats. Specification of these SV Sertoli cells is thought to be regulated by paracrine signals such as RSPO1, FGFs etc. from adjacent SOX17-positive RT cells (Imura-Kishi et al., 2021; Uchida et al., 2022; see review by Han et al., 2026). However, how SOX17-positive RT cells regulate SV formation in the proximal region of rodent testes remains unclear.

To address this question, we generated a novel transgenic (Tg) mouse line that exhibits excess SV formation by the Sertoli cells ectopically expressing *Sox17* under the control of the human *AMH* promoter (SOX17^+^ Sertoli cells). Such SOX17^+^ Sertoli cells enhance the expression of *WNT4* and *RSPO1* in SV and RT regions, respectively. Moreover, we showed a partial rescue by the SOX17^+^ Sertoli cells in the Sertoli valve formation and aberrant spermiogenesis in the cKO testes with *Sox17*-null RT. These findings highlight the importance of SOX17-downstream paracrine actions in the SV and RT for the proper valve formation, rather than the SOX17 roles in the RT formation in itself.

## Results

### Overexpression of SOX17 in Sertoli cells induced hyperplasia of the SV region

To investigate the downstream paracrine effects of *Sox17*, we generated *AMH-Sox17* transgenic (Tg) mice that ectopically express *Sox17* in Sertoli cells under the control of the human anti-Müllerian hormone (AMH) promoter (Fig. 1A, S2). Among three Tg founders, one F0 Tg male (#27) exhibited infertility and was smaller in size of testes compared to his wild-type littermates (Fig. 1B). Histologically, the most prominent abnormalities were confined to the terminal region of the seminiferous tubules, whereas the overall architecture and spermatogenesis of the remaining seminiferous tubules were largely preserved (Fig. 1C). In this Tg male, the Sertoli valve (SV) structure appeared to be expanded in the terminal segments of the seminiferous tubules (Fig. 1B, C). Additionally, abnormal aggregates of Sertoli cells were frequently observed on the luminal side of some seminiferous tubules near the RT (indicated by asterisks in Fig. 1B). Immunostaining for SOX17 and E-cadherin in the #27 Tg testis revealed ectopic and uniform expression of SOX17 in Sertoli cells throughout both the convoluted seminiferous tubules and the SV-like regions, similar to those observed in the RT epithelium (Fig. 1D). Extensive detachment of SV-like Sertoli cells was also observed near the RT, and the #27 F0 Tg male was infertile.

**Fig. 1.**
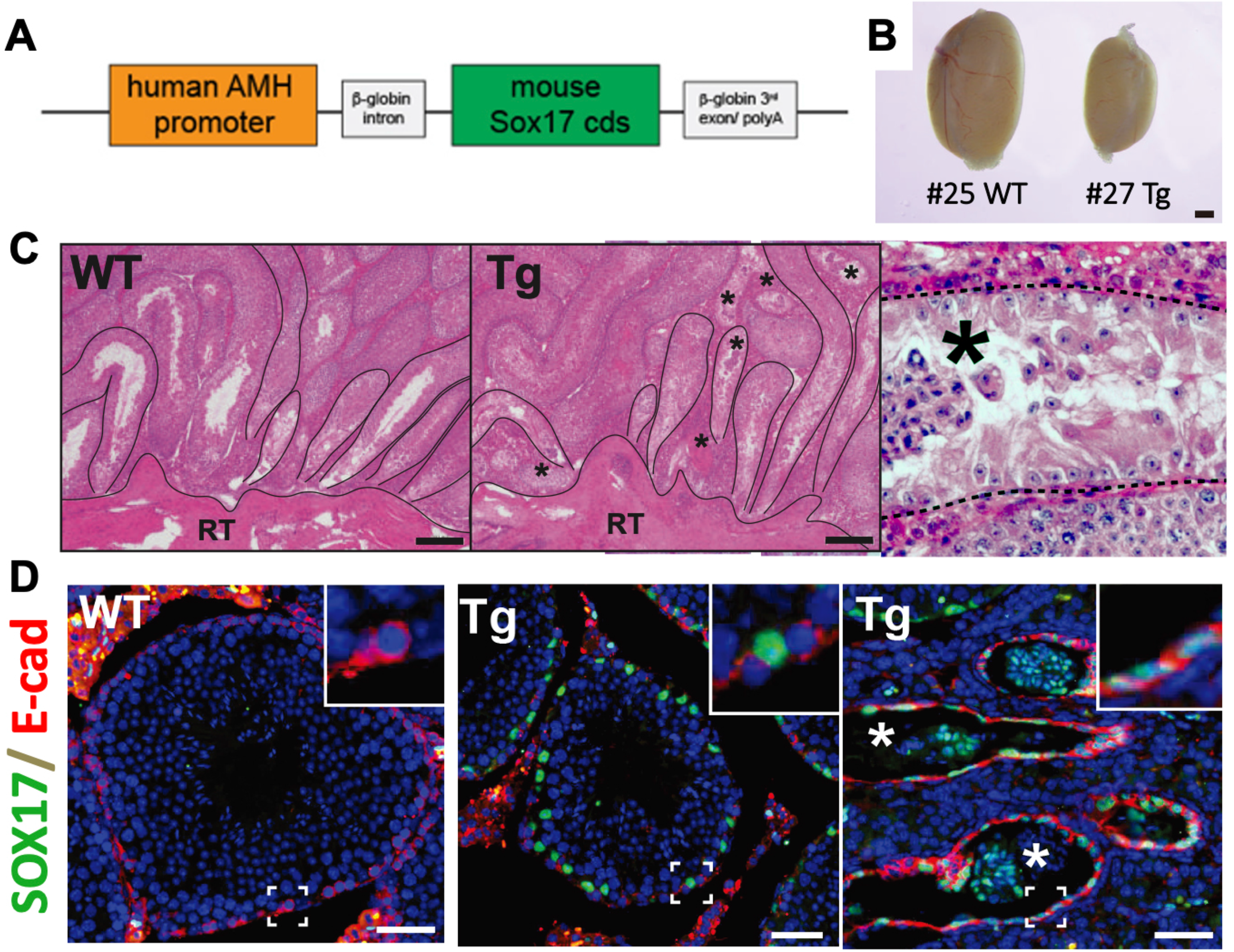
Hyperplastic SV structure in the *AMH-Sox17* Tg male mouse (F0) (**A**) Schematic diagram of the DNA construct containing *Sox17* coding sequences driven by the human *AMH* promoter. (**B**, **C**) Gross morphology and hematoxylin and eosin (H&E)-stained sections of testes from infertile Tg #27 and its wildtype littermates. In the Tg testis, normal spermatogenesis is partially observed in the convoluted seminiferous tubules (ST), while an expanded SV-like region including decidual cell aggregates is evident in the proximal area near the rete testis (RT) (asterisks in C). (**D**) Immunostaining for SOX17 and E-cadherin (E-cad; a marker for both RT epithelia and A_undiffer_ spermatogonia), showing ST regions (left two panels) and SV regions (rightmost panel) of the testis sections. The *AMH-Sox17* Tg testis exhibits ectopic expression of SOX17 in almost all of the Sertoli cells in both ST and SV. Asterisks in (D) indicate protruded SOX17-positive SV Sertoli cells within the E-cad-positive RT lumen. RT, rete testis; SV, Sertoli valve. Scale bars: 1 mm (B), 200 μm (C), 50 μm (D).

### Establishment of the AMH-Sox17 Tg Line (#26) with Excess SV Formation

One Tg male (#26) was fertile, and the *AMH-Sox17* transgene was successfully transmitted to its offspring (Fig. S2A, B). The morphology and weight of the testes in Tg mice were comparable to those of their wild-type littermates. Although the *AMH-Sox17* transgene was activated in a subset of Sertoli cells during both fetal and postnatal stages (Fig. S2C, D), an expansion of the acetylated tubulin-positive Sertoli valve (SV) region was observed in the proximal testis of this line (Fig. 2A), similar to that seen in the infertile #27 F0 Tg male with a more severe phenotype (Fig. 1D). Because this Tg line could be stably maintained by backcrossing beyond the 8th generation, line #26 was used for all subsequent experiments (hereafter referred to as “Tg”).

**Fig. 2.**
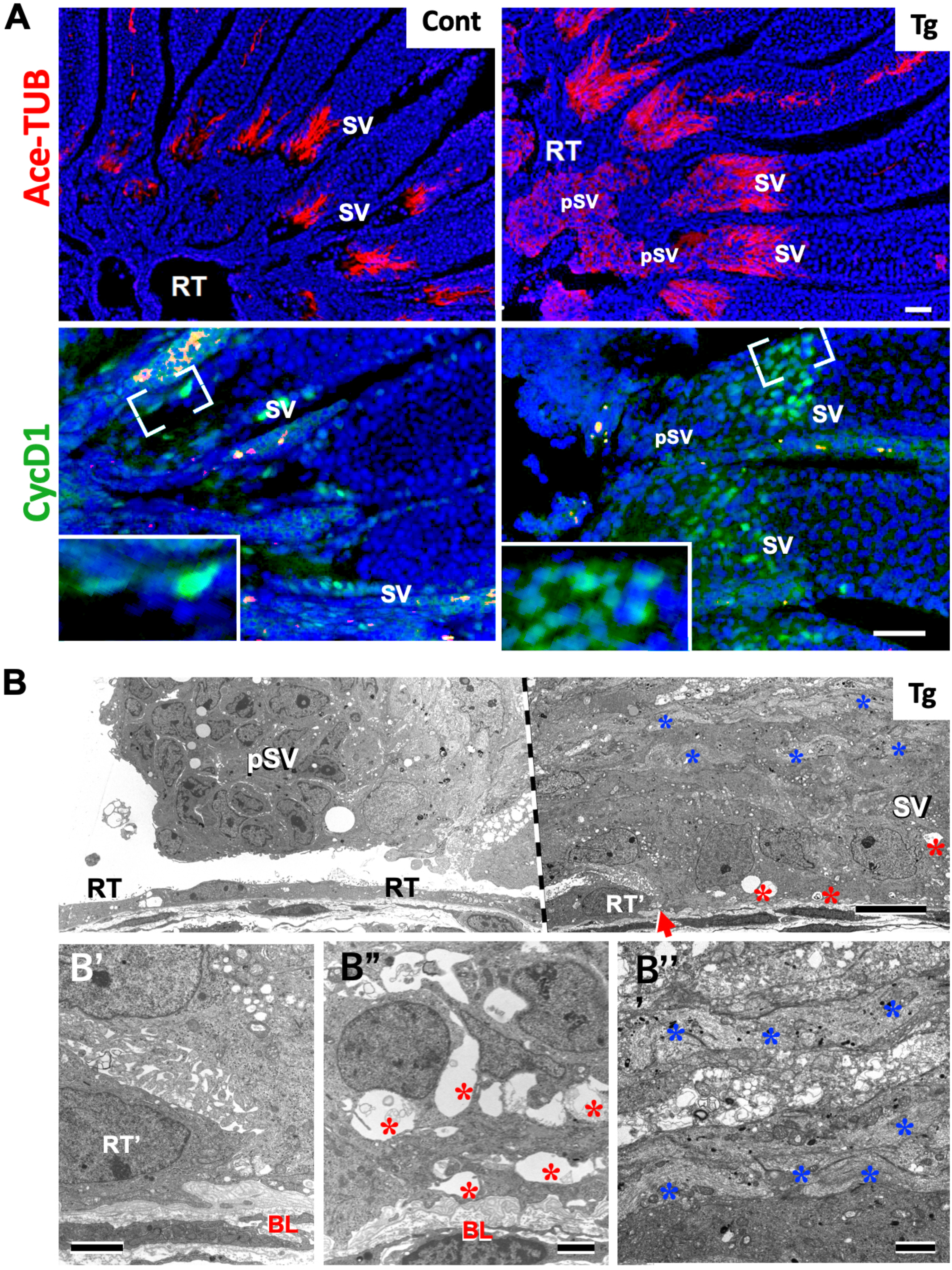
Morphological characterization of the hyperplastic SV structure in a novel *AMH-Sox17* Tg line. (**A**) Immunofluorescence staining for acetylated tubulin (red) and cyclin D1 (green) in the testes from Tg (#26) mice, showing expanded SV Sertoli cells. (**B**) Transmission electron microscopy (TEM) analysis of a Tg (#26) testis at the RT-SV boundary (red arrow), revealing abnormal features of SV Sertoli cells, such as protruded SV (pSV) structures extending into the RT lumen (upper left in B), multiple layers of cell processes containing microtubule bundles (blue asterisks), and open extracellular spaces in the basal region of the RT-SV boundary (red asterisk). Lower panels (B’–B’’’) are magnified views of the upper right panel in B. A vertical dashed line at the center marks the region hidden by the grid (grid spacing is 10 μm). BL, multilayered SV-typical basal lamina; RT, rete testis; SV, Sertoli valve. Scale bars: 50 μm (A), 10 μm (B), 2 μm (B’–B’’’).

In this Tg line, no appreciable alterations in the RT-SV region are observed during the immature stages up to 1 week after birth. However, following sexual maturation, immunostaining for acetylated tubulin and Cyclin D1 clearly marks an expanded SV region at the terminal segments of the seminiferous tubules (Fig. 2A). This expansion was further confirmed by electron microscopy, which revealed multiple cytoplasmic layers containing bundled microtubules extending distally from Sertoli cells into the rete testis (RT) lumen (blue asterisks, Fig. 2B). At the boundary between the SV and RT regions, SV Sertoli cells were supported by a multilayered basal lamina, consistent with previous reports (Fig. S3A; Tainosho et al., 2011; Uchida et al., 2022). Notably, in the Tg testes, the SV Sertoli cells displayed vacuole-like intercellular spaces in the basal portion of the terminal segment, and some appeared detached from the basal lamina (red asterisks, Fig. 2B). Furthermore, these cells were often found protruding into the RT lumen (designated as “pSV” in Fig. 2A, 2B, S3B), where necrotic or damaged cells were observed at the distal tips of the protrusions. These findings suggest that the protruding SV Sertoli cells may undergo necrosis and be sloughed off into the RT lumen.

### Partial rescue of defective spermatogenesis and infertility in RT-specific Sox17 cKO mice by AMH-Sox17 Tg

In RT-specific *Sox17* conditional knockout (cKO) testes, deformation of the Sertoli valve (SV) leads to backflow of rete testis (RT) fluid into the seminiferous tubules, resulting in defective spermiogenesis and male infertility (Uchida et al., 2022). In contrast, *AMH-Sox17* Tg mice exhibit excessive SV formation due to ectopic expression of SOX17 in SV Sertoli cells; however the *AMH-Sox17* transgene is not activated in the RT epithelium. To evaluate whether this ectopic expression could rescue the SV defects in the RT-specific *Sox17* cKO background, we examined the testis phenotype of double mutant mice (*Sox17* cKO; *AMH-Sox17* Tg, referred to as cKO;Tg) (Fig. 3). By crossing Tg; *Sox17^f/f^* males with *SF1-Cre;Sox17^f/f^*females, we generated four genotypes for comparison: control (*Sox17^f/f^*), cKO (*SF1-Cre;Sox17^f/f^*), Tg (*Tg; Sox17^f/f^*), and cKO;Tg (*Sf1-Cre;Sox17^f/f^; Tg*).

**Fig. 3.**
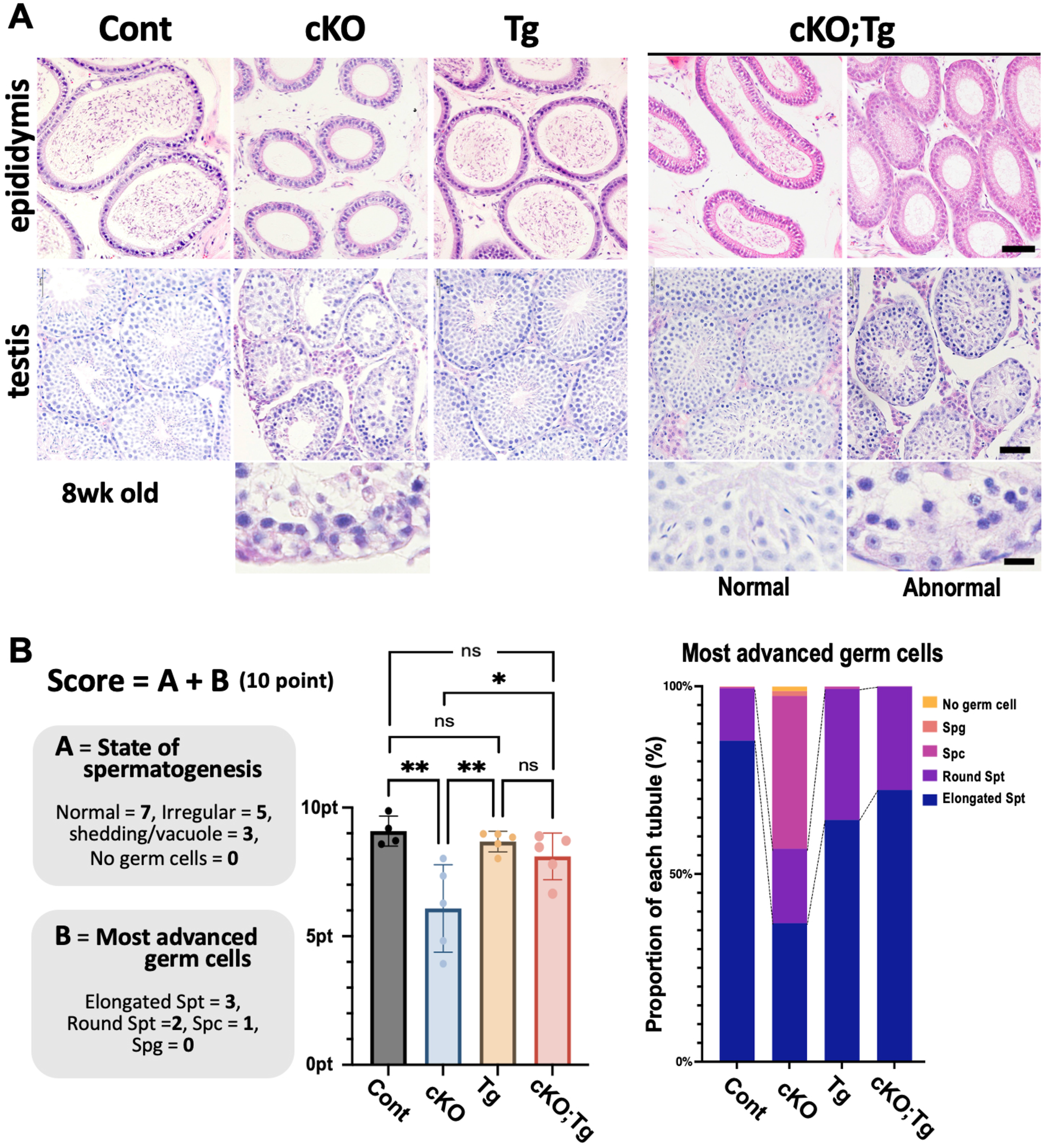
Morphometric analysis of spermatogenic quality in control (*Sox17^f/f^*), cKO (*Sf1-Cre;Sox17^f/f^*), Tg (*Tg; Sox17^f/f^*), and cKO;Tg (*Sf1-Cre;Sox17^f/f^; Tg*) testes. (**A**) H&E-stained sections of the epididymis and testes from control, cKO, Tg, and cKO;Tg males. Two representative testis images from both fertile and infertile cKO;Tg males are also presented. The lower panels present higher magnification images of damaged seminiferous epithelia. (**B**) Evaluation of spermatogenesis in cross-sections of convoluted seminiferous tubules (STs). The evaluation score (maximum 10 points) was calculated as the sum of the seminiferous tubule structural integrity score (Group A; 7 points) and the most advanced germ cell type score (Group B; 3 points). The two graphs on the right show the evaluation scores for each genotype (left; mean ± SD; *p<0.05; **p<0.01; one-way ANOVA) and the percentage of seminiferous tubules containing the most advanced germ cells (right). Defective spermatogenesis observed in cKO testes was rescued in the cKO;Tg double mutant testes. Scale bars: 50 μm (top two rows); 10 μm (bottom row) in A.

As expected, control and Tg males showed normal spermatogenesis and the presence of spermatozoa in the epididymis. In contrast, cKO males exhibited abnormal spermiogenesis with a complete absence of epididymal spermatozoa (Fig. 3A), consistent with previous findings (Uchida et al., 2022). Interestingly, cKO;Tg double mutants showed improved spermatogenesis, with the presence of epididymal spermatozoa and a partial restoration of fertility compared to cKO males. Quantitative assessment of spermatogenesis using a 10-point scoring system (7 points for seminiferous tubule integrity + 3 points for presence of advanced germ cells) confirmed a significant improvement in cKO;Tg males (Fig. 3B) related to cKO males. This was further supported by an increased ratio of elongated spermatids in these cKO;Tg testes. Notably, epididymal spermatozoa were detected in 2 of 7 cKO;Tg individuals at 8 weeks of age. Furthermore, to assess fertility, prolonged cohabitation mating trials were performed using a separate group of four cKO;Tg males. Two of the four males successfully sired offspring, although these animals were not the same as the cohorts used for the initial and quantitative evaluation of spermatogenesis.

These results demonstrate that ectopic SOX17 expression in Sertoli cells partially rescues the structural and functional defects of the SV caused by the absence of SOX17 in the RT, thereby improving spermatogenesis and restoring fertility in otherwise infertile males.

### Ectopic SOX17 expression in Sertoli cells rescues SV deformation in RT-specific Sox17 cKO testes

In RT-specific *Sox17* conditional knockout (cKO) testes, disruption of the SV formation leads to defective spermiogenesis and male infertility (Uchida et al., 2022), which could be partially rescued by the ectopic SOX17 expression in Sertoli cells, as described above (Fig. 3). To assess whether such ectopic SOX17 expression could rescue the SV defects caused by *Sox17* deficiency in the RT, we examined the RT-SV phenotypes of *Sox17* cKO;*AMH-Sox17* Tg double mutant (cKO;Tg) mice (Fig. 4).

**Fig. 4.**
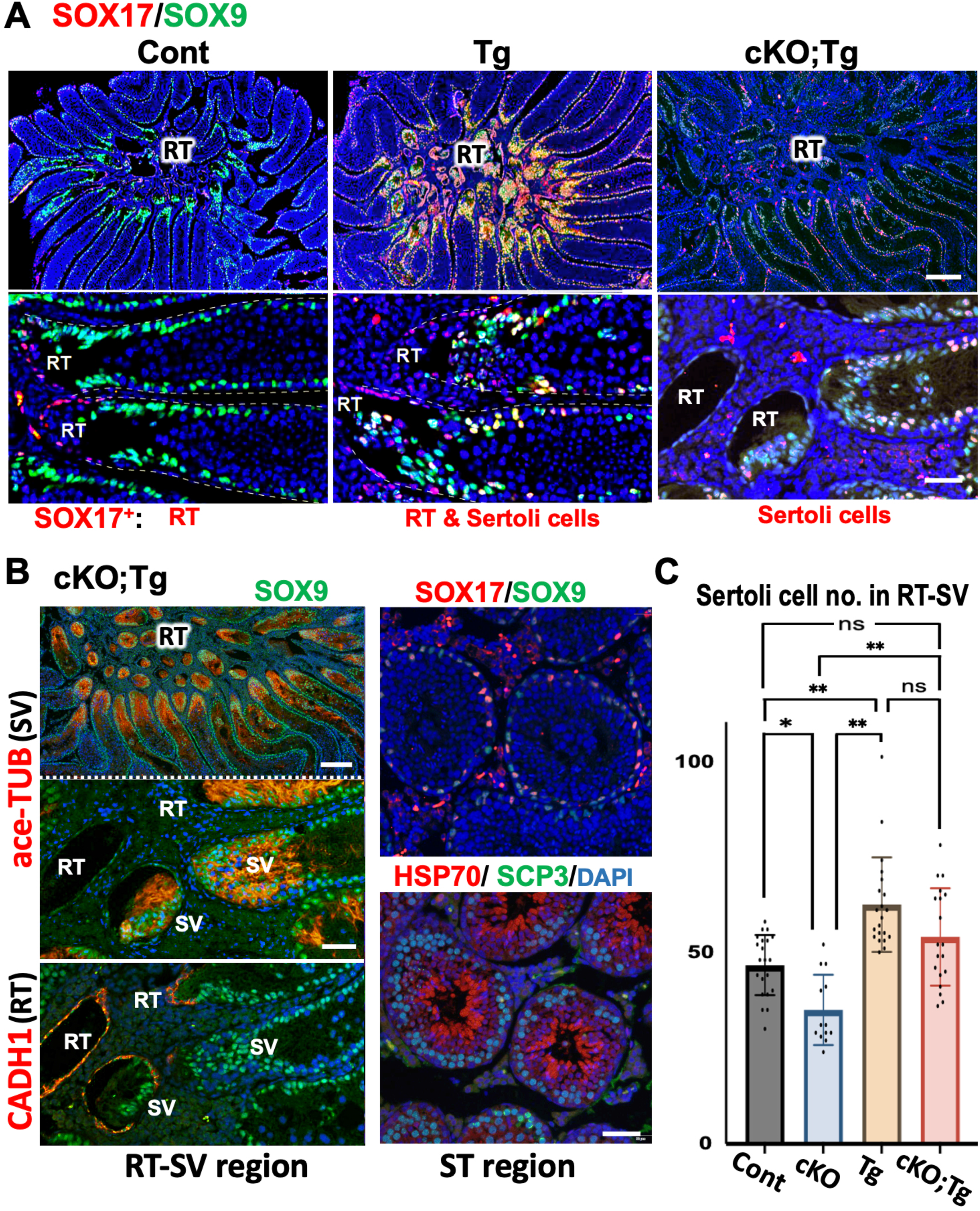
Partial normalization of the expanded SV structure in *AMH-Sox17* Tg testes by cKO;Tg double mutants. (**A**) Immunostaining for SOX9 and SOX17 in the RT–SV–ST region of proximal testes in control, Tg, and cKO;Tg mice. Ectopic SOX17 expression is observed in Sertoli cells of Tg testes, whereas SOX17 expression is completely absent in RT cells of the cKO;Tg double mutants. The expanded region of SOX17/SOX9 double-positive SV Sertoli cells (yellow) seen in Tg testes appears to be reduced in cKO;Tg double mutants. (**B**) Immunostaining for SOX9 (green) and acetylated tubulin (SV marker; red) or E-cadherin (RT marker; red) demonstrates the restoration of proper SV formation in the RT–SV region of the cKO;Tg double mutants. Additional panels show immunostaining for SOX17 (red)/SOX9 (green) and HSP70 (elongated spermatid marker; red)/SCP3 (spermatocyte marker; green) in ST regions (left panels). (**C**) Quantitaive data of the nuclear density of Sertoli cells in the RT-SV region (mean ± SD; *p<0.05; **p<0.01; one-way ANOVA). RT, rete testis; SV, Sertoli valve. Scale bars: 200 μm (top row) and 50 μm (bottom row) in A and B.

Double immunostaining for SOX17 and SOX9 confirmed that in cKO;Tg testes, SOX17 expression was restricted to Sertoli cells in the terminal segments of the seminiferous tubules, and was absent from the RT epithelium—opposite to the expression pattern seen in control testes, where SOX17 was confined to RT epithelial cells (Fig. 4A). In Tg testes, both RT epithelial cells and SV Sertoli cells were SOX17-positive. An expanded region of SOX17/SOX9 double-positive SV Sertoli cells was evident in Tg testes, while this region appeared to be reduced in cKO;Tg testes, suggesting partial normalization of the SV region (yellow area, Fig. 4A).

Furthermore, acetylated tubulin-positive valve structures were properly formed in cKO;Tg testes, and normal spermatogenesis including SCP3-positive spermatocyte and Hsp70-positive elongated spermatids was observed in the convoluted seminiferous tubules (Fig. 4B). Quantitative analysis of SV Sertoli cell density revealed that the number of SOX9-positive nuclei in the RT-SV region was significantly increased in Sox17 Tg testes and significantly decreased in *Sox17* cKO testes compared to control testes (Fig. 4C). Interestingly, in the cKO;Tg double mutant testes, SV Sertoli cell density was significantly higher than that in cKO testes, showing an intermediate level between Tg and cKO testes, comparable to that of controls (Fig. 4C). These findings indicate that ectopic SOX17 activity in SV Sertoli cells can rescue the structural defects of the SV in RT-specific *Sox17* cKO testes, in a *Sox17* dose-dependent manner.

Previous studies have shown that *Rspo1* is expressed in RT epithelial cells in wild-type mice, but its expression is markedly reduced in cKO testes (Uchida et al., 2022). Along with the known expression of *Wnt4* in SV Sertoli cells, we finally examined the expression patterns of *Rspo1* and *Wnt4* in the RT-SV region across different genotypes using RNAscope in situ hybridization (Fig. 5). Fixation, sectioning, and in situ hybridization were performed under identical conditions on serial sections. Expression profiles of RT-specific *Rspo1* and SV-specific *Wnt4* were generally comparable among control, Tg, cKO, and cKO;Tg mice (Fig. 5A), which confirm no ectopic *Rspo1* expression in SOX17^+^ SV Sertoli cells. In Tg testes, *Wnt4* expression appeared to be enhanced in the SV region, consistent with the expanded SV phenotype in these mice (Fig. 2). Notably, RT-specific *Rspo1* expression was increased in Tg testes and slightly elevated in cKO;Tg mice relative to cKO mice, suggesting a potential paracrine mechanism, in which ectopic SOX17 activity in Sertoli cells might influence *Rspo1* expression in adjacent RT epithelial cells.

**Fig. 5.**
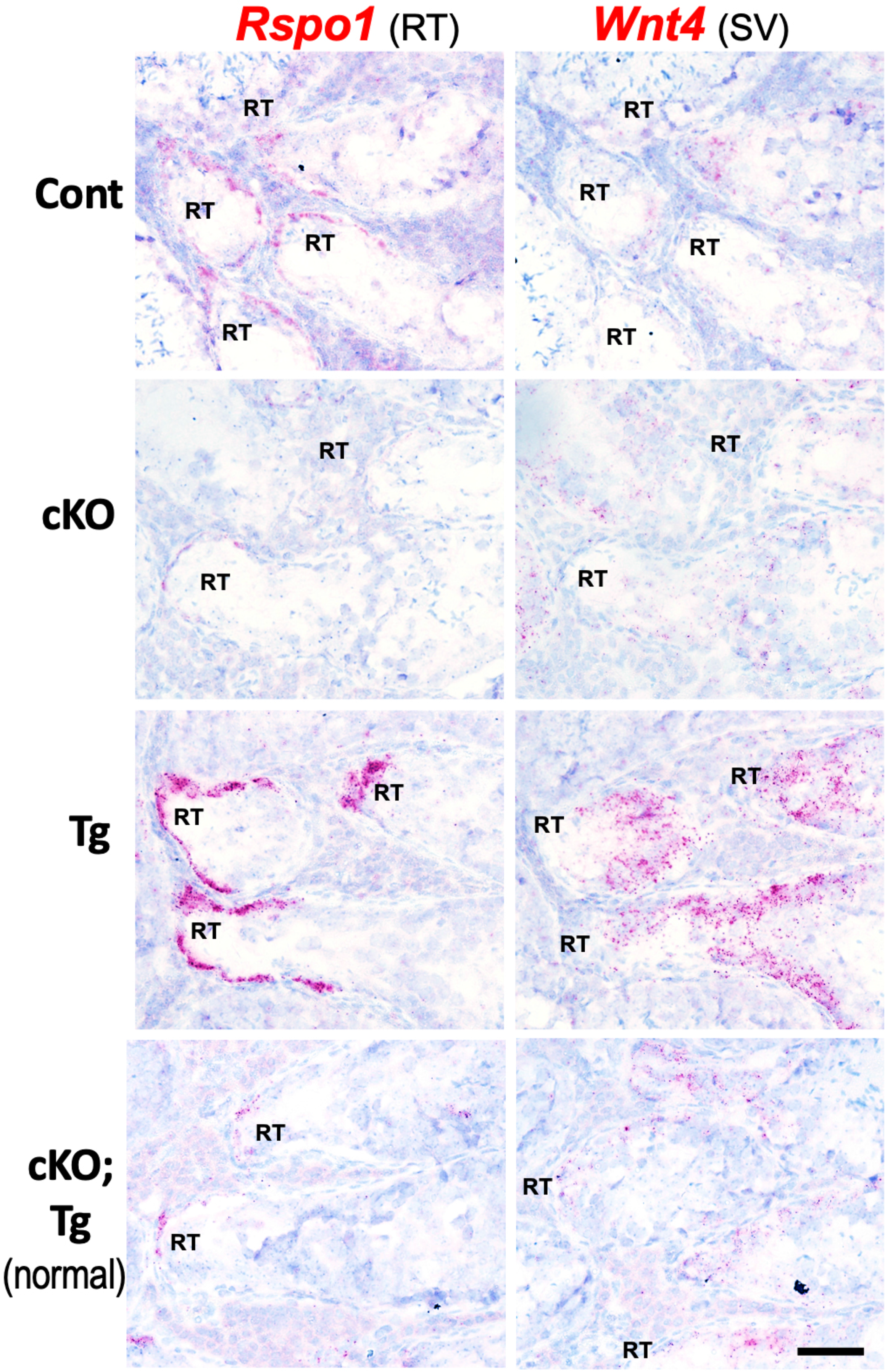
Enhanced *Rspo1* and *Wnt4* expression in the RT-SV boundary of the testes in *AMH-Sox17* Tg mice. RNA *in situ* hybridization showing the *Rspo1* and *Wnt4* expression profiles of RT-SV boundary in four genotypes: control (*Sox17^f/f^*), cKO (*Sf1-Cre;Sox17^f/f^*), Tg (*Tg; Sox17^f/f^*), and cKO;Tg (*Sf1-Cre;Sox17^f/f^; Tg*) testes. RT, rete testis; SV, Sertoli valve. Scale bar, 50μm.

## Discussion

A previous study using RT-specific *Sox17* cKO mice demonstrated that *Sox17* expression is critical for Sertoli valve (SV) formation and proper spermiogenesis. Deletion of *Sox17* in the rete testis (RT) disrupted the specification of modified Sertoli cells that form the SV, leading to backflow of RT fluid and abnormal detachment of immature spermatids within the seminiferous tubules (Uchida et al., 2022; Fig. 6; Fig. S1A). These findings established the essential role of SOX17⁺ RT epithelia in SV formation and clarified the physiological function of the SV in modulating fluid dynamics within the seminiferous tubules in vivo.

**Fig. 6.**
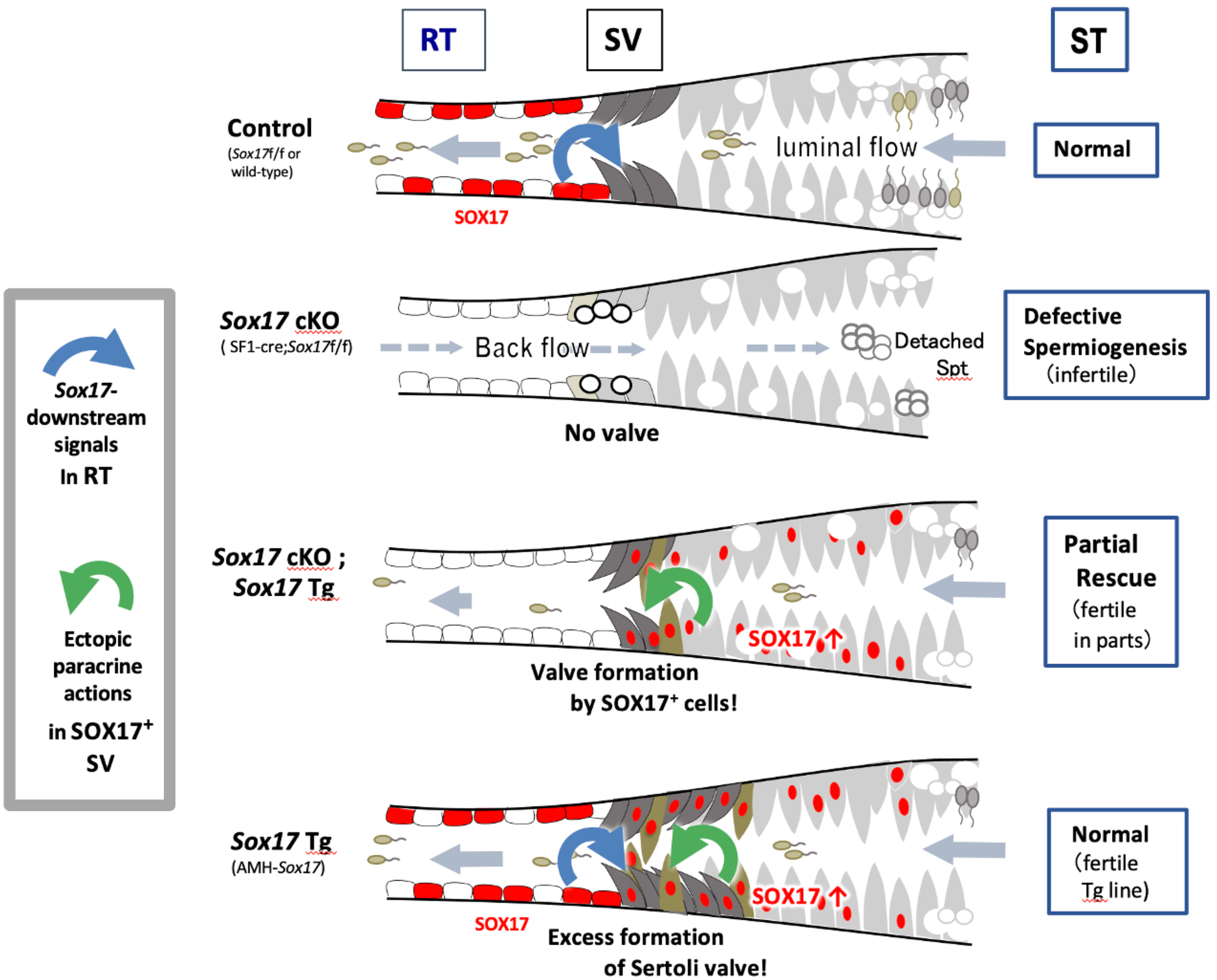
Schematic representation of valve structure size regulation by the *AMH–Sox17* Tg mouse line. RT-specific *Sox17* cKO mice exhibit deformed Sertoli valve (SV) structures, resulting in defective spermiogenesis and male infertility (Fig. 3; Uchida et al., 2022). Ectopic expression of SOX17 in Sertoli cells induces excessive SV formation in the terminal segments of the seminiferous tubule (ST), along with elevated Rspo1 expression in the rete testis (RT). Notably, this ectopic SOX17 expression partially rescues epididymal sperm production and restores fertility in cKO;Tg males. These *Sox17* Tg mice, displaying SV hyperplasia and its functional rescue, provide valuable models for investigating paracrine signaling mechanisms within the RT-SV region.

In the present study, we demonstrated that ectopic expression of SOX17 in Sertoli cells induces excessive SV formation in the seminiferous tubules, along with increased *Rspo1* expression in the RT. Moreover, this ectopic SOX17 expression partially rescued epididymal sperm production and restored fertility in cKO;Tg males (Fig. 6). These results suggest that SOX17 activity in the RT can be, at least in part, functionally substituted by its ectopic expression in Sertoli cells. The observed fertility recovery in cKO;Tg males, even in the absence of SOX17 in RT cells, implies that the primary role of SOX17 in the RT is restricted to SV formation in the terminal segments and is not essential for sperm transport and maturation in the RT-to-epididymis pathway in mice.

Ectopic SOX17 expression in Sertoli cells may mimic paracrine signaling normally mediated by SOX17⁺ RT epithelia in the SV region. One possible explanation is that RT epithelial cells and Sertoli cells share lineage similarities (Malolina and Kulibin, 2019; Uchida et al., 2022; Sato et al., 2026). Several key transcription factors of Sertoli cells — including WT1, NR5A1, SOX9, GATA4, and, in part, DMRT1— are also expressed in the RT epithelia (Aiyama et al., 2015; Malolina and Kulibin, 2019; Kulibin and Malolina, 2022; Uchida et al., 2022; Malolina et al., 2023). During sexual maturation, ectopic SOX17 expression in Sertoli cells may induce paracrine and/or autocrine signaling programs resembling those of RT cells, thereby promoting hyperplasia in Tg testes and facilitating SV recovery in cKO;Tg testes.

The *Sox17* transgene is ectopically activated in Sertoli cells not only near the SV but also throughout the testis, whereas the SV phenotype is restricted to the RT-adjacent region of Tg testes (Figs. 1C, 2A, 4A). One possible explanation for this spatial discrepancy is that SOX17 may act on RT-lineage Sertoli-like cells located in the terminal segments adjacent to the RT. A recent study by Mayère et al. (2022) reported that PAX8⁺ RT progenitor cells, referred to as early supporting-like cells, contribute to the proximal region of the seminiferous tubules, suggesting heterogeneity in the origin of Sertoli cells near the RT during embryogenesis, even though PAX8 expression in fetal Sertoli cells is downregulated shortly after testis differentiation. It is therefore possible that ectopic SOX17 expression in these rare supporting-like cells may induce paracrine signaling similar to that of RT epithelia, thereby leading to SV phenotypes in Sertoli cells near the RT in Tg mice.

Otherwise, SOX17 has been shown to directly modulate genes encoding cell adhesion and extracellular matrix components, and to broadly influence basement membrane-associated gene networks during tubular morphogenesis (Niimi et al., 2004; Niakan et al., 2010; Walters et al., 2023). SOX17 may enhance the responsiveness of SOX17⁺ Sertoli cells to RT-derived morphogenetic signals through the establishment of an RT-like extracellular matrix microenvironment, thereby promoting spatially restricted SV formation near the RT during postnatal maturation. Indeed, RT epithelia produce high levels of growth factors and cytokines, including FGFs, TGFs, and WNT signals (Imura-Kishi et al., 2020; Uchida et al., 2022).

Interestingly, *Rspo1* and *Wnt4* signaling is activated in the RT-SV region (Uchida et al., 2022; this study). In *Wnt4*-null embryos, the RT region is underdeveloped at the seminiferous tubule–RT junction, leading to a failure in connecting the RT and seminiferous tubules, similar to the rete ovary (RO) phenotype (Mayère et al., 2022). These findings suggest that *Wnt4–Rspo1* signaling is essential for proper connection of the terminal segment of seminiferous tubules to the RT and efferent ducts, which is critical for the integrity of the male reproductive tract. Our in situ hybridization analyses showed enhanced *Rspo1* and *Wnt4* expression driven by SOX17⁺ Sertoli cells in Tg mice (Fig. 4), which likely contributed to the rescue of SV formation observed in cKO;Tg mice (Fig. 5). The presence of protruding SV cells expressing Cyclin D1 supports the idea that SOX17-mediated paracrine activity may involve localized *Rspo1-Wnt4* signaling during SV formation. Considering the potential involvement of other signaling pathways such as FGFs and TGFs, further studies are warranted to delineate the specific roles of individual paracrine factors in the RT–SV region prior to SV formation. Moreover, the present study raises several important questions regarding how SV Sertoli cells adjacent to the RT region may influence the total number of Sertoli cells per testis, and how RT-lineage Sertoli-like cells contribute to the support of spermatogenesis in the vicinity of the SV. Addressing these issues will be essential to determine how RT-derived signaling, fluid dynamics, cellular composition, lineage heterogeneity, and RT architecture collectively regulate SV formation at the RT–SV interface.

We acknowledge several limitations in the present study. First, quantitative molecular analysis of the SV region remains technically challenging. The Sertoli valve is a very small transitional structure, with approximately 20 sites per mouse testis, and is confined to a narrow region within ∼100 μm from the terminal end of the rete testis-associated seminiferous tubules (Aiyama et al., 2015; Nakata et al., 2015; Imura-Kishi et al., 2020). Because of this limited spatial extent, selective isolation of SV tissue without contamination from adjacent regions, as well as acquisition of sufficient material for quantitative analyses such as qPCR, is currently difficult (Imura-Kishi et al., 2020). In addition, the highly interdigitated architecture of Sertoli cells, which are tightly connected via specialized junctional complexes, further complicates their dissociation into intact single-cell populations after sexual maturation. For these technical reasons, SV-associated molecular changes in this study had to be primarily evaluated using histological and immunohistochemical approaches. Second, while our previous and present studies support a role for SOX17 in regulating SV formation, the direct downstream target genes and molecular mechanisms remain unidentified, despite extensive scRNA-seq analyses of SV and RT cells in RT-specific *Sox17* cKO testes (Uchida et al., 2022). Using this *Sox17* Tg mouse line, future comprehensive transcriptomic approaches (e.g., bulk RNA-seq and spatial transcriptomics of the SOX17^+^ Sertoli cells) will make it possible to identify candidate downstream effectors and further validate the proposed model in which SOX17 regulates SV formation via paracrine signaling mechanisms.

In conclusion, by generating a novel *AMH-Sox17* Tg mouse line, we demonstrate that the size of the SV structure in the terminal segment of the seminiferous tubules can be experimentally manipulated. Furthermore, SV formation was successfully rescued in *Sox17* cKO testes by introducing the *AMH-Sox17* transgene into RT-specific *Sox17* cKO mice. These genetically engineered models — exhibiting SV overdevelopment or restoration — provide valuable tools for investigating paracrine signaling and elucidating the biological significance of the SV and RT in the male reproductive system of mammals.

## Materials and Methods

### Animals

*AMH-Sox17* Tg mice were generated by pronuclear injection of the plasmid-based transgene into the C57BL6 background, as described previously (Shinomura et al., 2014). The transgene was prepared by inserting the entire coding region of the *Sox17* gene (Kanai-Azuma et al., 2002) downstream of the *hAMH* promoter-driven cassette (Fig. 1A). We obtained three independent F0 Tg males (lines #13, 26, and 27). Among the three founders, only line #26 was successfully established as a breeding line while maintaining spermatogenic capacity and was therefore used for most analyses in this study. Immunohistochemical analysis of Tg26 testes revealed mosaic, heterogeneous SOX17 expression among individual Sertoli cells. The #27 F0 male was infertile and exhibited extensive detachment of SV-like Sertoli cells that ectopically and uniformly expressed SOX17. Consequently, this #27 line could not be established as a breeding line. The RT-specific *Sox17* cKO mice (i.e., *SF1-Cre*;*Sox17*^flox/flox^ mice; Dhillon et al., 2006; Kim et al., 2007) were produced from the Jackson Laboratory (Uchida et al., 2022). The *SF1-Cre*:*Sox17*^flox/flox^;*AMH-Sox17* Tg double mutant (cKO;Tg), *AMH-Sox17* Tg;*Sox17* ^flox/flox^ (Tg), *SF1-Cre*;*Sox17*^flox/flox^ (cKO) and *Sox17* ^flox/flox^ (control) mice were mainly generated by crossing the *AMH-Sox17* Tg; *Sox17* ^flox/flox^ males with *SF1-Cre*:*Sox17* ^flox/flox^ females. These virgin mutant males at 4 and 8 weeks of age were used for the subsequent phenotypic analyses. Note that a separate cohort was utilized for this mating trials, distinct from the animals subjected to histopathological analyses of spermatogenesis. To maximize the analysis of the anatomically minute RT–SV region, the testes and epididymides from these virgin males were dissected into the proximal quarter containing the RT–SV region (for SV analysis) and the remaining tissue blocks for each animal (Figure S4). Consequently, due to these technical requirements, tissue allocation strategy, and independent evaluation of distinct histological parameters, the exact numbers of animals analyzed differed across individual assays.

All animal experiments were performed following the Guidelines for Animal Use and Experimentation at the University of Tokyo, and all experimental procedures performed herein were approved by the Institutional Animal Care and Use Committee, Graduate School of Agricultural and Life Sciences, University of Tokyo (approval IDs: P13-762, P13-764, P22-147). The PCR list for the genotyping was shown in Table S1.

### Histology and immunohistochemistry

Testes were fixed in 4% PFA at 4°C overnight, dehydrated through an ethanol series, and embedded in paraffin. For the quantitative analysis of the RT-SV region, testes were transected after fixation and surface dye marking at approximately one-quarter of their length from the rete testis (RT) side. The quarter containing the RT and the proximal seminiferous tubules was used for analysis of the Sertoli valve (SV) region, whereas the remaining three-quarters were used to evaluate spermatogenesis. During the dehydration process, the RT-containing tissue block, whose cutting surface was pre-labeled with hematoxylin, was placed and gently pressed beneath a coverslip (Fig. S4), flattening the specimen and allowing the seminiferous tubules to spread radially from the RT while orienting the RT–SV–seminiferous tubule (ST) axis nearly parallel to the sectioning plane. The tissue was subsequently embedded in paraffin while maintaining this orientation. Serial sections were cut parallel to the flattened surface, enabling frequent acquisition of sagittal RT–SV–ST profiles (Fig. 4A; S4).

The paraffin sections (4 μm thickness) were subjected to hematoxylin and eosin (H&E) staining, PNA lectin (acrosome) staining and immunohistochemistry. For immunohistochemistry, the sections were incubated overnight at 4°C with the primary antibodies listed in Table S2. Immunoreactions were visualized using Alexa fluor-488/555/594/680-conjugated secondary antibodies (Abcam). Samples were analyzed by fluorescence microscopy (BX51N-34-FL-2) or Leica TCS SP8 confocal laser microscopy. Testes from at least four animals in each group were examined, and tissue sections from cKO, Tg and cKO;Tg and their littermate controls were processed in parallel to ensure the reproducibility of the results.

### Electron microscopy

Testes were trimmed and fixed in 2.5% glutaraldehyde in 0.1 M sodium cacodylate buffer (pH 7.4) overnight, post-fixed with 0.5% OsO4 suspended in 0.1 M phosphate buffer (pH 7.3) for 30 min, and dehydrated through a graded ethanol series. After passing through propylene oxide, the tissues were embedded in Epon 812. Ultrathin sections were cut, stained with uranyl acetate and lead citrate, and observed by TEM (JEM-1010; JEOL).

### RNA in situ hybridization

Testes were fixed in 10% formalin, dehydrated through an ethanol series, and embedded in paraffin. The paraffin sections (4 μm thickness) were processed for RNA in situ detection using the RNAscope 2.5HDAssay Red Manual (ACDBio) as described previously (Imura-Kishi et al., 2021; Uchida et al., 2022). The probes used in this study are listed in Table S3.

### Evaluation of Spermatogenesis score

To evaluate spermatogenesis in mouse testes, we performed the fluorescence staining with PNA lectin (a marker for acrosome; Nakata et al., 2015) on tissue sections (Control: n = 4; cKO: n = 4; Tg: n = 5; cKO;Tg: n = 5 males). Fluorescent images of entire testis sections were acquired using a confocal microscope (TCS SP8 Leica Microsystems) with the Tiling function in LAS X software. A two-axis scoring system was applied: Group A categorized the morphological condition of the seminiferous tubules, and Group B assessed the most advanced germ cell stage present within each tubule. Scores were assigned as maximun 10 points as follows: Group A evaluated the morphological condition of the seminiferous tubules, while Group B assessed the most advanced germ cell stage present in each tubule. Scores were assigned with a maximum of 10 points (Group A [tubule condition]: normal [7 points]; disrupted spermatocyte stage [5 points]; partial cell dropout or loss [3 points]; Sertoli cells only [0 points]) plus Group B [most advanced germ cell type]: elongated spermatids [3 points]; round spermatids [2 points]; spermatocytes [1 point]; spermatogonia only [0 points]) (see Fig. 3B).

### Quantitative morphometric analyses

The number of Sertoli cells in the RT-SV region was quantified using the RT-SV-ST spreading sections immunostained for SOX9, E-cadherin, and DAPI. Only sagittal sections showing a continuous RT–SV–ST axis and sectioning the SV approximately through its mid-sagittal plane were included in the quantitative analysis. Based on the distribution of spermatogonial stem cells, the ∼100 μm region extending from the RT boundary along the seminiferous tubule toward the ST side was defined as the SV region (Aiyama et al., 2015). The SV region was operationally defined as comprising (i) the terminal 100 μm segment of the seminiferous tubule immediately adjacent to the rete testis (RT) and (ii) the protruded SV extending into the RT lumen (Fig. 4C; S4). All SOX9-positive/E-cadherin-negative Sertoli cell nuclei within this region were counted.

### Statistical analysis

For comparisons among four genotype groups, one-way ANOVA followed by Tukey’s multiple comparisons test was performed. Quantitative data are presented as means ± standard deviation (SD) and shown as bar graphs with dot plots. Statistical analysis was performed using GraphPad Prism (v.10.4.2; GraphPad Software, San Diego, CA, USA). *P* < 0.05 was considered indicative of significance, and levels of significance are shown as \**P* < 0.05 and \*\**P* < 0.01. In addition, single comparisons between two groups, as shown in Fig. S2A, were were analyzed using a two-tailed Student’s t-test (data are presented as means ± standard error of the mean).

## Acknowledgments

The authors wish to thank Y. Hirate, H.M. Takase, and Y. Kuroda for their technical advice and support; Ikuno Oike for her secretarial assistance. The authors acknowledge Y. Akimoto, S. Matsubara, and J. Hayakawa for their excellent TEM technical support.

## Author contributions

X.H., A.U., K.N., T.E., A.U., M. K-A., Y.K. designed the study. X.H., A.U., S.L., K.N., K.T., A.Y., M. K-A., A.K., and R.H. performed the biological experiments. The manuscript was written by X.H., A.U., S.L., and Y.K.

