## Supplementary material for "Ectopic *hAMH*-driven SOX17 expression induces hyperplastic Sertoli valve formation in mouse testes": sup figs1-4 & tables S1-3

### Supplementary Figures

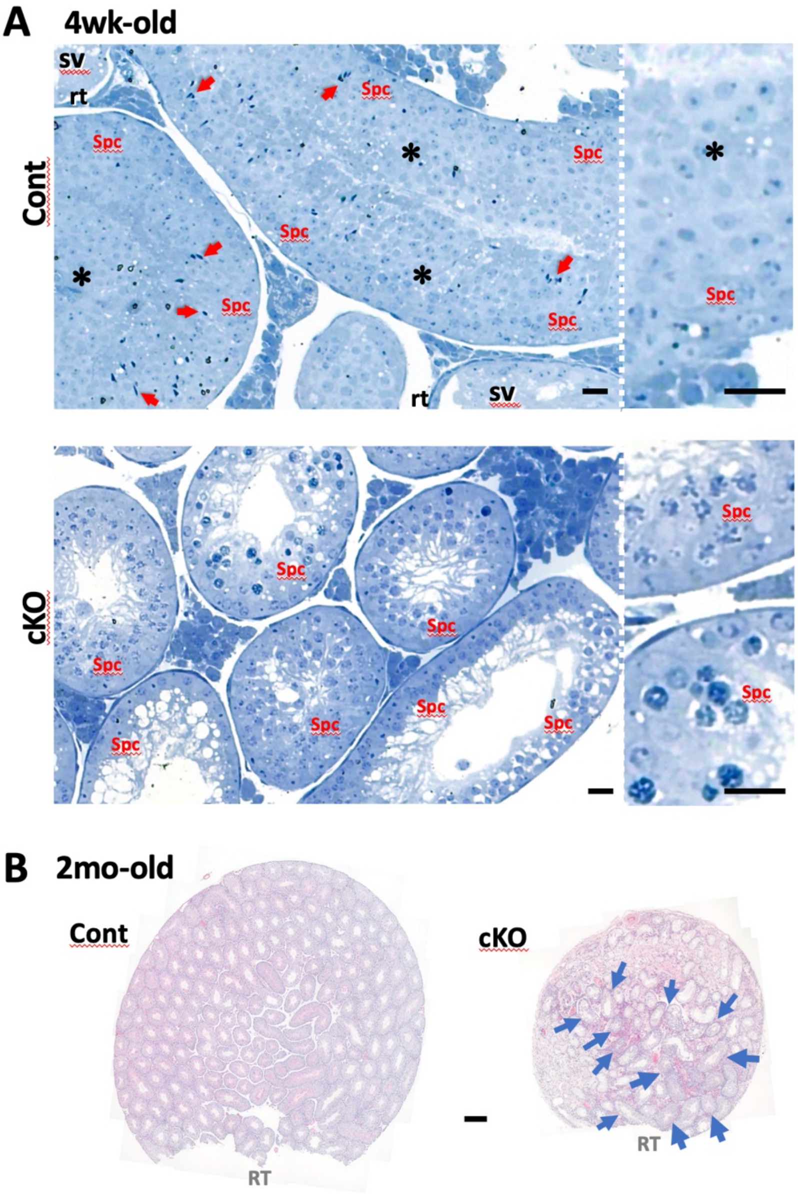


**Figure S1. Early and later phenotypes of immature and mature testes in the control (*Sox17*$^{f/f}$) and *Sox17* cKO (*Sf1-Cre;Sox17*$^{f/f}$) mice**

(**A**) Early phenotype of the convoluted seminiferous tubules adjacent to the rete testis (RT)–Sertoli valve (SV) region in a 4-week-old *Sox17* cKO testis (toluidine blue-stained Epon sections), showing preservation of the Sertoli cell processes at the sites of spermatid detachment. Round (asterisk) and elongated (red arrows) spermatids are observed in the control testis, not in the *Sox17* cKO testis. (**B**) Late phenotype of the whole testis (after dissection of the RT–SV region) from approximately 2-month-old control and *Sox17* cKO littermates (H&E-stained paraffin sections). The severity of spermatogenic defects varies among individual seminiferous tubules, forming a mosaic pattern of progressive degeneration resembling the reported distribution of individual seminiferous tubular loops (Nakata et al., 2015). Blue arrows indicate seminiferous tubules with relatively mild defects located in close spatial proximity to one another. RT, rete testis; Spc, spermatocyte; SV, Sertoli valve. Scale bars: 20 µm (A); 200 µm (B).

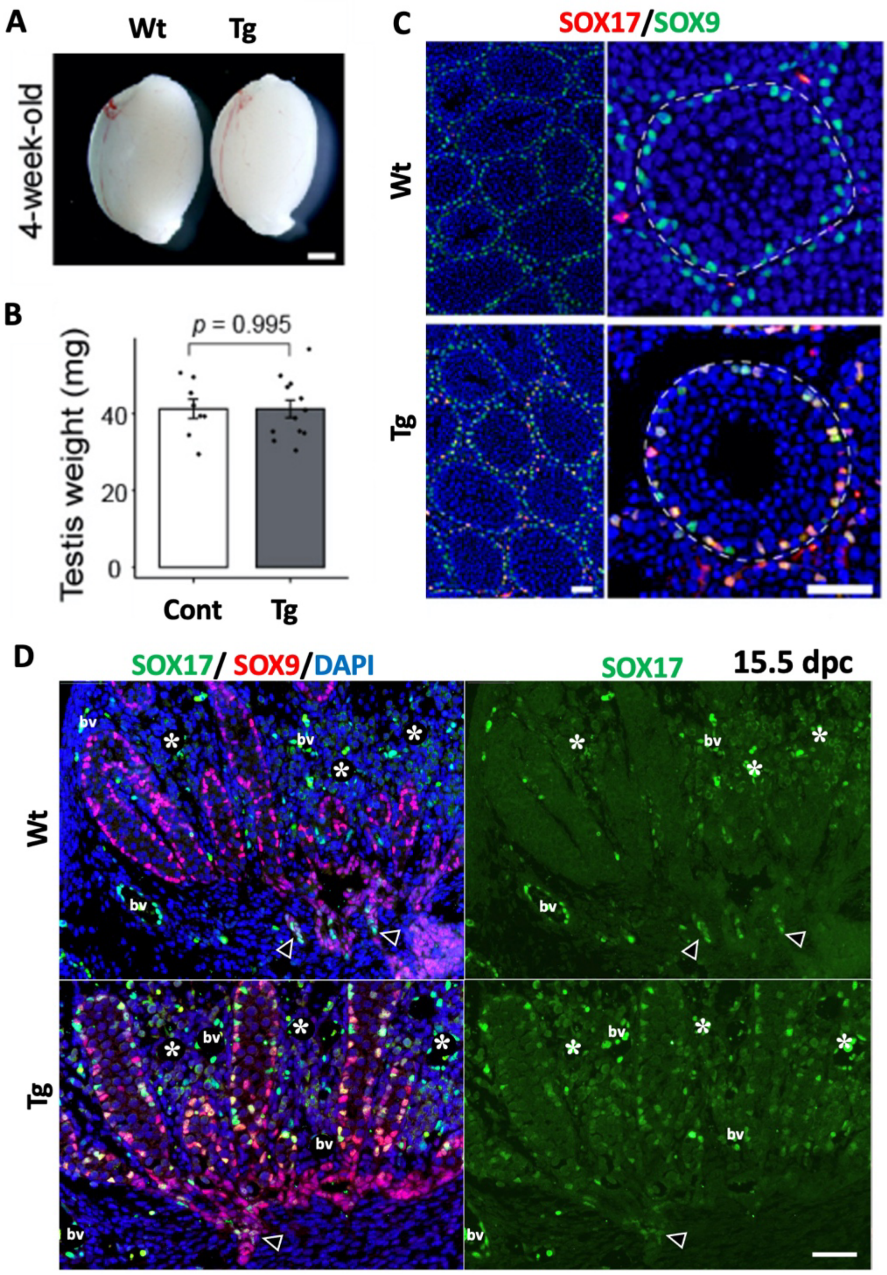


**Figure S2. Establishment of the *AMH-Sox17* Tg Line (#26) with moderate ectopic expression of SOX17**

(**A**, **B**) The gross morphology (A) and weight (B; means ± standard error) of the testes from the Tg (#26) and wildtype littermates. (**C**) Immunofluorescence double staining for SOX17 (red) and SOX9 (green) in Tg (#26) testes, showing ectopic SOX17 expression in Sertoli cells (yellow in merged image). (**D**) Immunofluorescence double staining for SOX17 (green) and SOX9 (red), with DAPI nuclear counterstaining (blue), in Tg (#26) and wild-type embryos at 15.5 days post-coitum (dpc). SOX17-positive signals are detected in fetal Sertoli cells of Tg testis, but not in wild-type one. Arrowheads indicate SOX17-positive fetal RT cells. Asterisks indicate the testicular interstitium. Bv, blood vessel. Scale bar, 1mm (A), 50 µm (C, D).

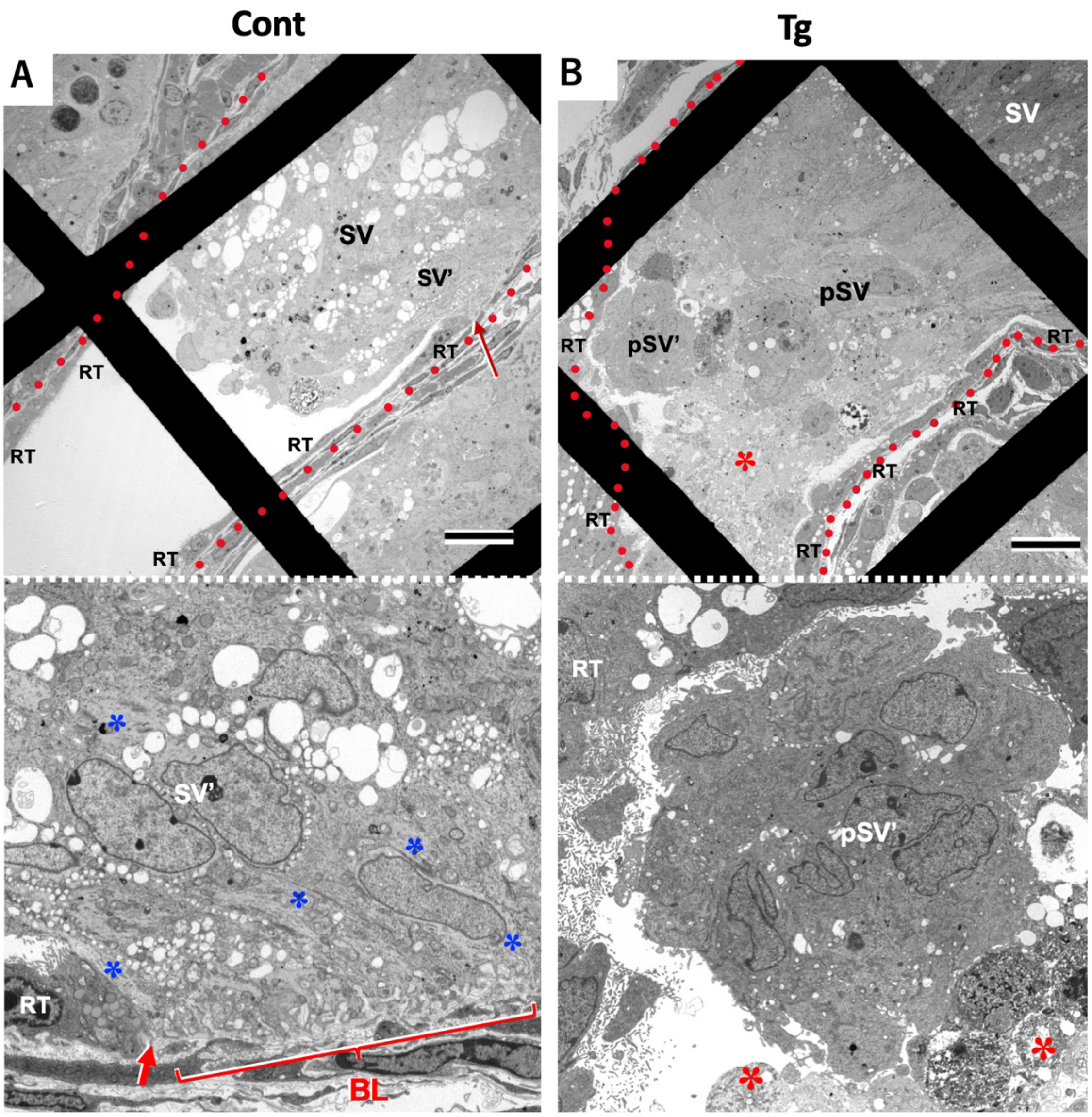


**Figure S3. Transmission electron microscopy (EM) analysis comparing the Sertoli valve (SV) between wild-type and transgenic (Tg) testes**

(**A**) TEM image of a wild-type (control) testis at the SV–rete testis (RT) boundary. A multilayered basal lamina (BL) and prominent microtubule bundles (blue asterisks) are observed in the SV Sertoli cells adjacent to the RT epithelium (red arrow, the RT-SV border). (**B**) TEM image of a Tg (#26) testis at the RT-SV boundary, showing protruded SV Sertoli cells (pSV) and their necrotic cells (red asterisks) within the RT lumen. The pSV is located quite far from the original SV structure (SV), which is shown in the upper right corner. Lower panels show higher magnification of the SV’ and pSV’ of upper panels. Red circle dots indicate the out line of the RT on the EM grid. BL, multilayered SV-typical basal lamina; RT, rete testis; SV, Sertoli valve. Scale bar, 20μm.

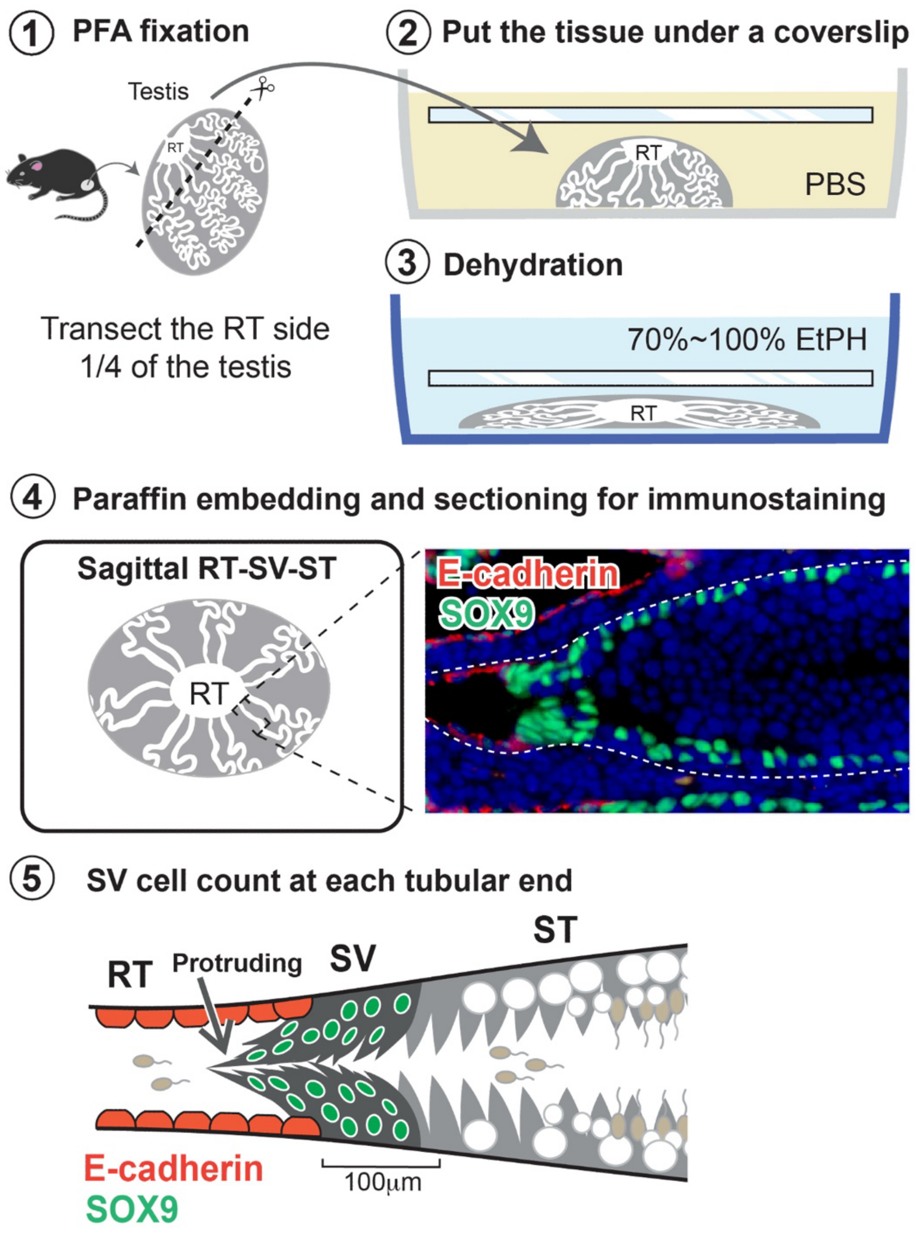


**Figure S4. Procedure for morphometric analysis of the SV–RT–ST region of the testis**
**Step 1.** Approximately one-quarter of the testis on the rete testis (RT) side was excised after PFA fixation. **Steps 2-3.** The RT-containing tissue was placed beneath a coverslip, followed by dehydration through 70% and 100% ethanol, allowing the seminiferous tubules to spread radially from the RT and orienting the RT–SV–ST axis nearly parallel to the sectioning plane. This procedure enabled frequent acquisition of sagittal RT–SV–ST profiles. **Step 4.** The flattened tissue was embedded in paraffin and serially sectioned parallel to the flattened surface, followed by immunostaining for SOX9 (Sertoli cell marker, green), E-cadherin (rete testis marker, red), and DAPI (blue). **Step 5.** SV cells located within 100 μm of the RT–SV border, including protruding SV Sertoli cells extending into the RT lumen, were counted for each tubular end.

**Supplementary Table S1** / List of primers used for mouse genotyping

| Primer Name | Sequence(5'→3') |
|---|---|
| Cre-F | ATTTGCCTGCATTACCGGTC |
| Cre-R | ATCAACGTTTTCTTTTCGGA |
| Sox17-flox Fw2 | AGATGTCTGGAGGTGCTGCTCACTGTAACG |
| Sox17-flox Rev | TGCCTCTTTTGCCGAACACACAAAAGGAGC |
| Ex4-Sox17-Fw2 | CCTCGGGGATGTAAAGGTGAA |
| Ex5-Sox17-Rev2 | GGTCAACGCCTTCCAAGACT |

**Supplementary Table S2** / List of antibodies used in this study

| Antigen | Dilution | Description | Company | CAT# |
|---|---|---|---|---|
| Ace-TUB | 1/400 | Mouse monoclonal | Sigma | T6793 |
| E-cad | 1/200 | Mouse monoclonal | BD Transduciton lab | 610181 |
| DDX4 (MVH) | 1/1000 | Rabbit polyclonal | Abcam | ab13840 |
| SCP3 | 1/100 | Mouse monoclonal | Santa Cruz | sc74569 |
| SOX9 | 1/200 | Rabbit polyclonal | Merk Millipore | ab5535 |
| SOX17 | 1/200 | Goat polyclonal | R&D systems | AF1924 |

**Supplementary Table S3** / List of probes used in this study

| Gene symbol | CAT# | Accession No. |
|---|---|---|
| *Rspo1* | 401991 | NM_138683.2 |
| *Wnt4* | 401109 | NM_009523.2 |
